# Single cell sequencing reveals microglia induced angiogenesis by specific subsets of endothelial cells following spinal cord injury

**DOI:** 10.1101/2022.01.25.477640

**Authors:** Chun Yao, Yuqi Cao, Yehua Lv, Dong Wang, Yan Liu, Xiaosong Gu, Yongjun Wang, Xuhua Wang, Bin Yu

## Abstract

Spinal cord injury (SCI) results in dynamic alterations of the microenvironment at the lesion site, which inevitably leads to neuron degeneration and functional deficits. The prominent deterioration of the milieu, derived from the destruction of spinal vascular system, not only activates innate immunity but also makes cells in the lesion lose nutrient supports. Limited endogenous angiogenesis happens after SCI, but the cell events at the lesion site underpinning this process have not been delineated so far. Here, we conducted single-cell RNA sequencing (scRNA-seq) of tissues in the spinal lesion at different time points after rat SCI. After performing clustering and cell-type identification, we focused on the vascular endothelial cells (ECs), which play a pivot role in angiogenesis, and drew a comprehensive cellular and molecular atlas for endogenous angiogenesis after SCI. We found that microglia and macrophage promote endogenous angiogenesis by regulating EC subsets through SPP1 and IGF1 signal pathways. Our results indicated that immune cells promotes angiogenesis by the regulation of specific cell subsets of vascular ECs, which provides new clues for the development of interventions for SCI.

## Introduction

Spinal cord injury (SCI) results in a complex physiological and pathological process that worsens functional outcomes (Hutson and Di Giovanni, 2019). Different from peripheral nervous system (PNS), the regenerative capacity of central nervous system (CNS) after an injury is limited (Tran et al., 2018). Multiple factors have contributed to the failure of CNS regeneration, including the low intrinsic growth capacity of axons, the existence of myelin-associated inhibitors, formation of the glial scar, and deteriorated microenvironment (Tran et al., 2018). Among those factors, the injury-induced microenvironment of CNS has overwhelming biological importance, as it undergoes dynamic composition of detrimental and beneficial factors from early to homeostatic stages, which profoundly affects neuropathology (Nishimura et al., 2013; Fan et al., 2018). Thus, to create a regenerative microenvironment for functional recovery, it is prerequired to clarify the molecular and cellular processes associated with the detrimental or beneficial factors following SCI.

Under normal physiological conditions, the vascular system can transport oxygen, nutrients, and hormones. Also, it carries away metabolic wastes, promotes cell circulation, and plays a pivotal role in maintaining elaborate functions of the nervous system (Haggerty et al., 2018). The SCI initially disrupts the vascular network in the injury microenvironment. Also, the permeability of the blood-spinal barrier will be changed, leading to ischemia, inflammatory response, and further tissue damage. These complex alterations in the vascular system hinder the regeneration and functional recovery of the damaged axons (Sweeney et al., 2019; Jin et al., 2021). Meantime, reconstruction of the vascular network in the injury microenvironment occurs. Reduction of the vascular endothelial cells (ECs) in the injured microenvironment within 24 hours after SCI results in decreased vascular density and therefore, benefits the local angiogenesis and vascular remodeling (Oudega, 2012; Halder et al., 2018). The endogenous angiogenesis occurs at the 3-4th day after SCI and emerges apparent vascular tissue on the 7th day (Bakshi et al., 2004). However, the density of new blood vessels around the lesion area is low, and the neovessels lack interaction and communications with other cell types, suggesting that the premature network of new blood vessels is unable to support the repair of injured spinal cord (Ng et al., 2011; Oudega, 2012).

Single-cell RNA sequencing (scRNA-seq) has recently been utilized to analyze the spinal cord tissues under developing (Delile et al., 2019), adult (Blum et al., 2021), and pathological conditions (Milich et al., 2021). The cellular heterogeneity, new cell subsets, and their interactions have thereafter been identified. These investigations have provided novel clues on the adaptive and pathological reactions of neurons and glial cells challenged by SCI. However, little information is available about the states of vascular ECs in the lesion microenvironment after SCI, which participate in angiogenesis and the formation of the vascular network. It is known that vascular ECs are heterogeneous in the CNS comparing with those in other tissues (Feng et al., 2019) (Vanlandewijck et al., 2018). But, how the heterogeneous vascular ECs are pathologically regulated in the injured spinal cord remains elusive. Here, we performed a scRNA-seq study to identify the heterogeneity of ECs in the blood vessels following rat SCI, and further illustrated the interactions and signaling pathways between ECs and other cell types, which play key roles in angiogenesis after SCI.

## Results

### Blood Vessel Alterations Around the Lesion Area after SCI

SCI results in direct vascular damage and initiates a cascade of events that alter the permeability of the blood-spinal cord barrier (BSCB) (Rocha et al., 2018; Tran et al., 2018). Typically, ECs form close contact with microglia, astrocytes, and vascular mural cells (pericytes and vascular smooth muscle cells (vSMCs)) (Yamazaki et al., 2021). To investigate the interactions between ECs and other cells, a rat crash SCI model was constructed. The alterations of blood vessels and relevant cells were then detected at 0 d, 3 d, and 7 d after SCI. As shown in Figure 1, in the normal spinal cord, the blood vessels with different diameters were concentrated in the gray matter area and were rich in number and crisscross. At 3 days after injury, in the injured area, the blood vessels were with reduced number and irregular shape, and their continuity was also interrupted. As the lesion area spreaded rostrocaudally with a spindle shape, a cavity gradually formed at 7 days after SCI. Blood vessels were scattered around the cavity. After SCI, microglia (IBA1 marked), a type of immune cells, migrated to the injured area and were activated as characterized by de-ramified and shorten processes. We also found that in normal condition, very few microglia contacted with blood vessels. However, after injury, the EC-constituted blood vessels were surrounded with microglia, indicating that microglia might interact with ECs and play an important role in regulating angiogenesis after SCI. In addition, we also detected vascular smooth muscle cells (vSMCs) (αSMA marked) and astrocytes (GFAP marked) in the lesion after SCI, which also showed some interactions with ECs (Supplementary Fig. S1). Collectively, there was a significant reduction in blood vessel density after SCI and ECs have an interaction with other cell types in the lesion microenvironment.

**Fig. 1.**
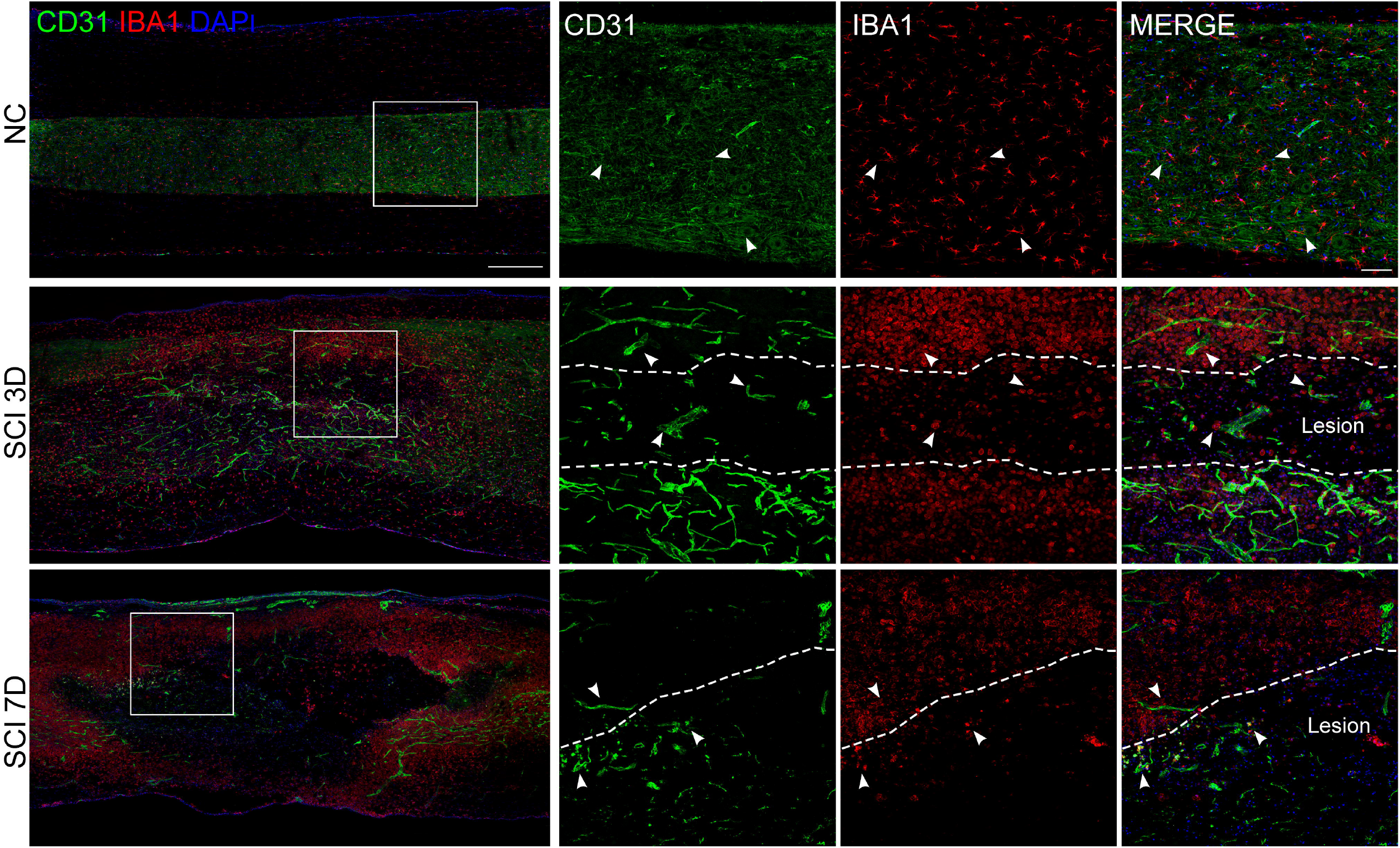
Immunostaining of ECs and microglia in the injured spinal cord tissues. Co-immunostaining of CD31^+^ECs (green) and IBA1^+^ microglia (red). Arrows indicate the interacted ECs and astrocyte. The three small squares in the right were the enlarged images of the white square in each rectangle image. The border between uninjured and injured was drawn with the dashed line. Nuclei were stained with DAPI (blue). Scale bar: 500 μm and 100 μm, respectively.

### Identification of Cellular Heterogeneity after Rat Crash SCI by Single-Cell Sequencing

To the explore cellular composition at the lesion site, we performed 10X Genomics scRNA-seq on unsorted cells of rat spinal lesions at 3 d (SCI-3D), 7d (SCI-7D) after SCI, or sham spinal cord (NC-3D and NC-7D) (Fig. 2a). Sequencing quality metrics were similar across samples, excluding the technical variation among samples (Supplementary Fig. S2a&b and Table S1). After initial quality control assessment and removal of low-quality cells, we obtained a transcriptomic dataset for 30,657 recovered cells, which underwent unsupervised clustering (Macosko et al., 2015) and visualized using dimensionality reduction algorithm t-SNE (Hinton, 2008; van der Maaten, 2014). Cell populations in four spinal cord samples were clustered into 26 clusters (Supplementary Fig. S2c). According to their gene profiles and canonical markers, we combined these clusters into 14 cell type clusters, including microglia (C0), endothelial cell (C1), macrophage (C2), neutrophil (C3), pericyte (C4), fibroblast (C5), vSMC (C6), astrocyte (C7), erythrocyte (C8), oligodendrocyte precursor cell (OPC) (C9), B cell (C10), monocyte (C11), oligodendrocyte (C12) and T cell (C13) (Fig. 2b). The proportions of ECs, pericytes, vSMCs and microglia decreased after SCI, while the proportion of macrophages increased (Fig. 2c). We annotated all clusters based on literature-curated marker genes and the distributions of classical marker genes in each cell type were shown in Fig. 2d. In addition, the expressions of the top three differentially expressed genes (DEGs) in each cell cluster were demonstrated in a heatmap and the violin plots (Fig. 2e and 2f, Supplementary Table S2).

**Fig. 2.**
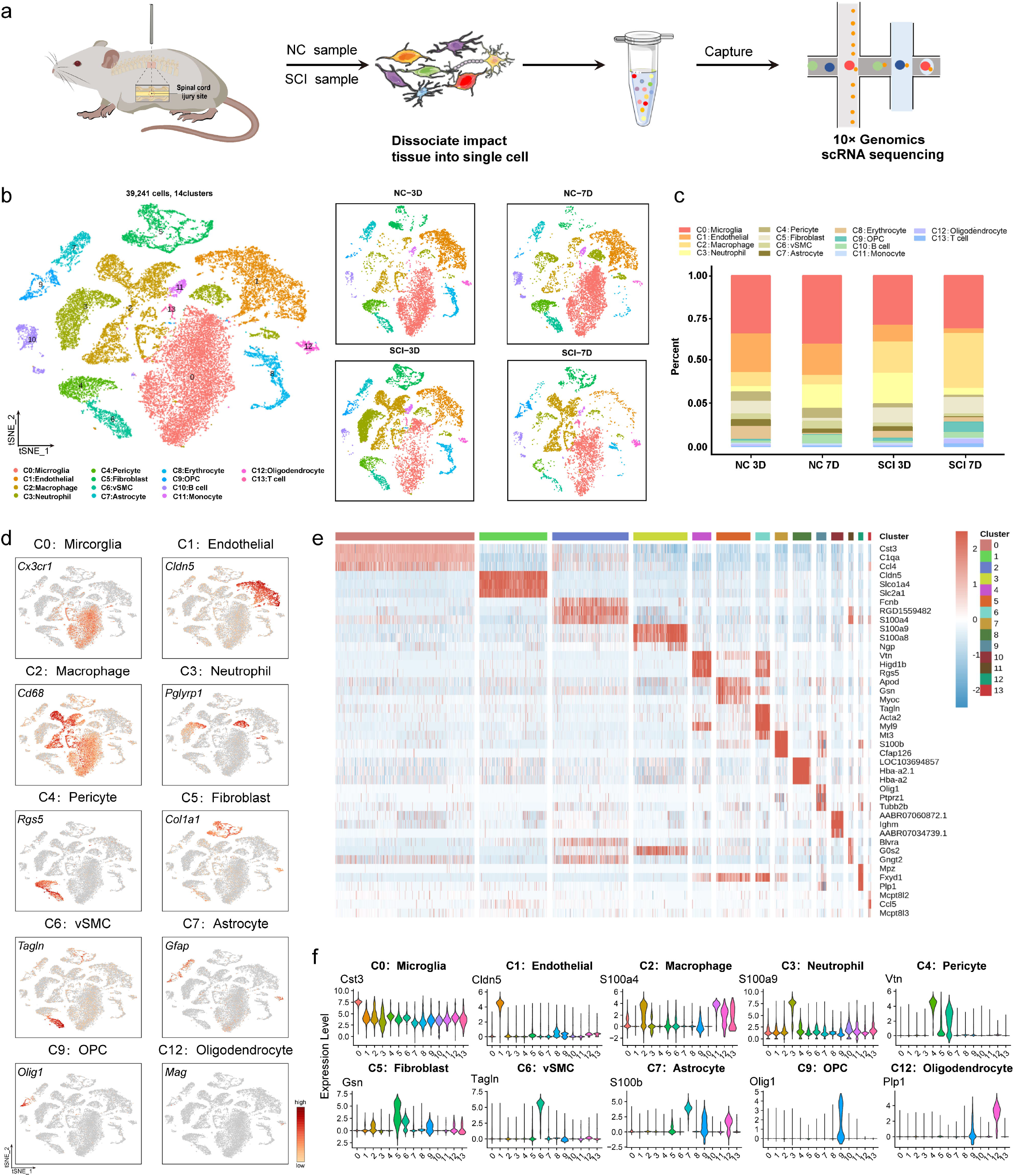
The single-cell sequencing landscape of spinal cord tissues after SCI. **a** Schematic of workflow for scRNA-seq using the 10x Chromium platform. **b** tSNE embedding of 37,325 cells annotated by cell types from jointly analyzed SCI and NC spinal cord tissues. Different cell clusters were color-coded. **c** The proportion of each cluster at different time samples. **d** tSNE maps showing the expression levels of the canonical biomarkers of each cell type, color-coded for the expression of marker genes. **e** The expression heatmap of the top 3 DEGs in tSNE-assigned cell populations. **f** Violin plots of the expression pattern of the DEG that best identifies each cell type.

### Cellular and Molecular Interactions between Different Cell Types after SCI

To systematically identify molecular interactions between cells in the spinal cord, we adapted CellPhoneDB (Efremova et al., 2020) (www.cellphonedb.org) to calculate “interaction scores” between ECs and other cell types. In each sample, a putative cell-cell communication network across cell types was constructed, in which nodes represented different cell types and the edges indicated the interactions between cell types. In the normal spinal cord, interactions were mainly enriched between EC, macrophage, microglia, and fibroblast. In contrast, distinct interactions were formed after SCI, within which the interactions between EC, fibroblast, and vSMC were highlighted (Fig. 3a). This analysis revealed considerable alterations in possible communication routes between ECs and other cell types. KEGG signaling pathway analyses were performed to identify pathways enriched in DEGs between ECs in different samples (SCI-3D vs NC-3D (Fig. 3b), SCI-7D vs NC-7D (Fig. 3c), and SCI-7D vs SCI-3D (Fig. 3d)). As shown, signaling pathways as focal adhesion, MAPK signaling pathway, ECM-receptor interactions were enriched in ECs after SCI. Fig. 3e and Supplementary Fig. S3 showed the detailed interactions between receptors on ECs and ligands from other cell types (Fig. 3e) or ligands from ECs and receptors on other cell types (Supplementary Fig. S3), respectively. Fig. 3f showed that in Fig. 3e, the receptors on ECs were mainly enriched in biological processes, including cell migration, regulation of chemotaxis, blood vessel development, and angiogenesis.

**Fig. 3.**
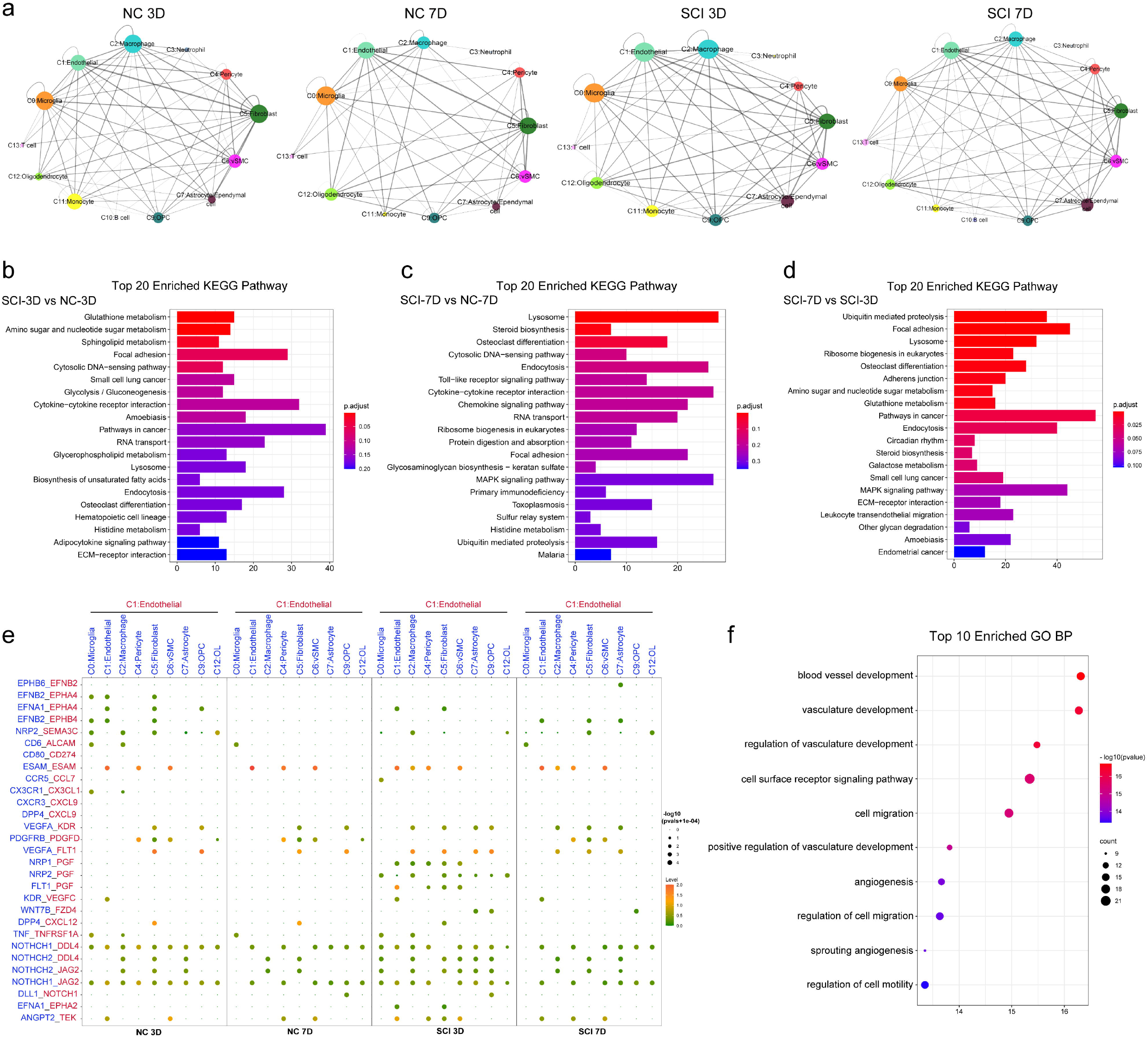
Cell-Cell Communication Networks in spinal cords after SCI Using CellPhoneDB. **a** Network visualization of ligand-receptor connectivity in NC and after SCI. Nodes represented clusters, the more interactions the cell with other cell types, the larger the node. Edges indicated the interactions between the nodes and the edges thickness of the edge was proportional to the number of receptor-ligand pairs between cell types. **b-d** Enriched top 20 KEGG pathways of differentially expressed genes in ECs between NC-3D and SCI-3D (**b**), NC-7D and SCI-7D (**c**), and SCI-3D and SCI-7D (**d**). The horizontal axis showed the gene numbers in each pathway, and the vertical axis showed the functional description of the enriched pathway. The color spectrum from blue to red represents the P-value. **e** Cell-type-specific interactions between ECs and the rest of the cell types when EC was the receptor cell. Circle sizes represented significance, which were defined as −log10 (p value+1e-04). Colors from green to red represented the interaction score between ligands and receptors. **f** Enriched top 10 GO BP terms of ECs receptor genes.

### Sub-Clustering of Endothelial Cells and Validation

Next, we further identified sub-types of ECs in the spinal cord. Overlaying canonical lineage markers onto unbiased clustering of 4,406 ECs, we clustered ECs into seven sub-clusters (EC0-6) (Fig. 4a). EC0, EC1, and EC2 were the major clusters, accounting for almost 75% of the total ECs, while the other four EC clusters (EC3, EC4 EC5, and EC6) constituted the resting (Fig. 4b&c). Notably, cell proportion analysis revealed that EC5 was almost absent in the uninjured spinal cord, but increased dramatically at 3 d, and decreased to normal by 7 d. The distributions of top DEGs of EC3 (Mt2A, Rnd1), EC5 (Angpt2, Apln), EC6 (Rgs5, Tagln), and other ECs were shown in violin plots (Fig. 4d and Supplementary Fig. S4a&b, Table S3). We then further classified sub-cluster identities according to previously reported typical EC subtype markers (Kalucka et al., 2020). The EC3, EC5, and EC6 were identified respectively as vein ECs, tip cells, and mural-like ECs based on their expression of the canonical marker (Mt2a, Angpt2, Rgs5). GO biological process (BP) enrichment analyses were performed of DEGs in each subset as shown in Fig. 4e and Supplementary Fig. S4c. As shown, the functions of DEGs in EC3 were enriched in cell proliferation, migration, and cell death. EC5, which was denoted as tip cells, was involved in angiogenesis, cell proliferation, and adhesion. This was consistent with previous reports (Hellstrom et al., 2007; Chen et al., 2019). EC6 was a set of cells that also expressed typical mural cell markers (Rgs5, Tagln). The GO enrichment results suggested that the EC subset might participate in the development of blood vessels. In addition, the pseudotime analysis of ECs was also performed. As shown in Fig. 4f, ECs were divided into two stages. Cells from sham control groups (NC-3D and NC-7D) were almost in stage 1, while cells from injured samples were in stage 2. As expected, EC0-EC4 and EC6 had a major distribution in stage 1, while EC5 distributed mainly in stage 2.

**Fig. 4.**
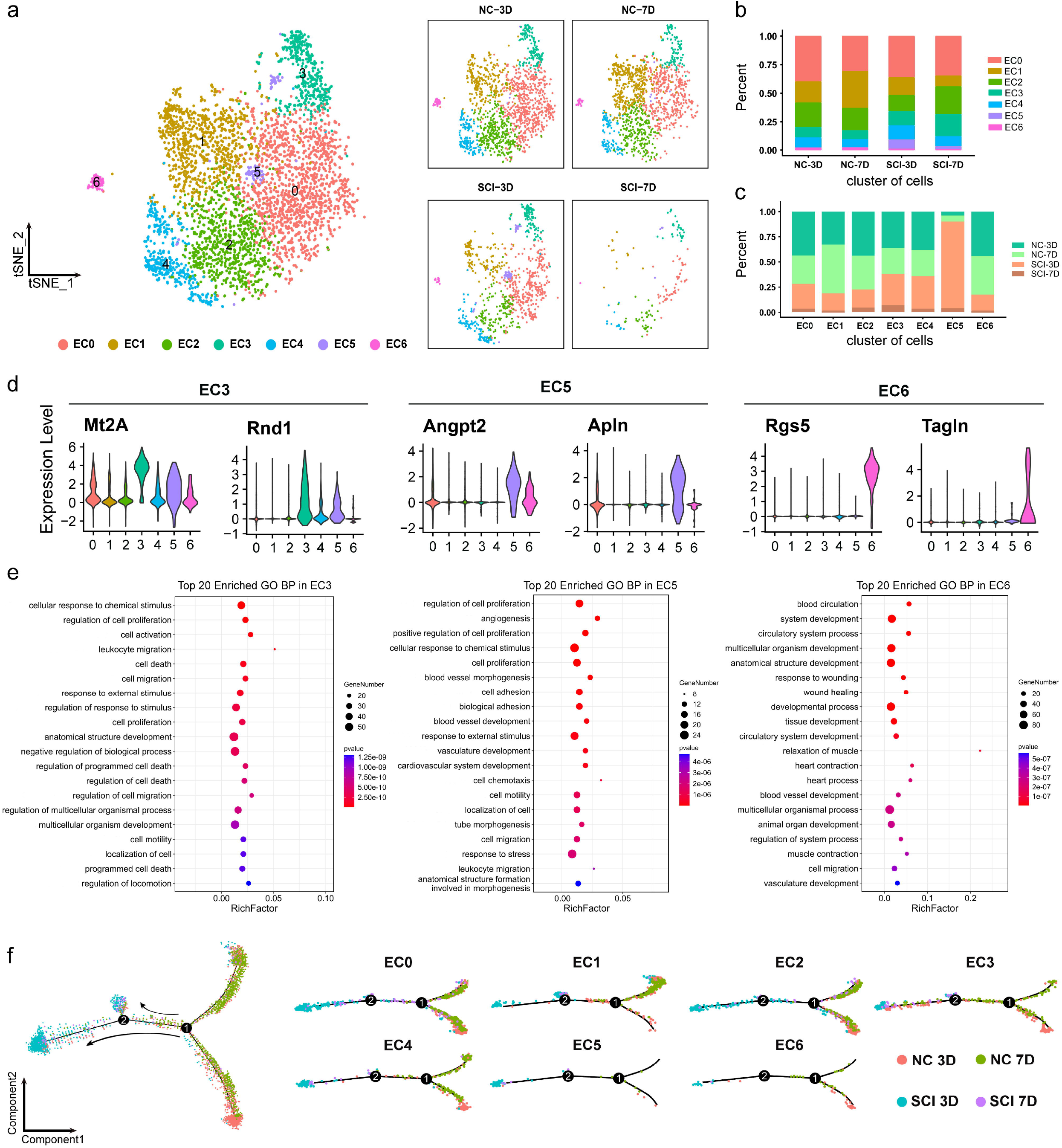
Molecular and temporal profile of ECs subtype heterogeneity after SCI. **a** tSNE maps of single-cell data for ECs (cell number n=7,140). Different cell clusters were color-coded. **b** The proportion of each EC sub-cluster at different time samples. **c** The percentage of different time samples cells in each EC sub-cluster. **d** Violin plots showed the differential expression distribution of two representative marker genes for EC3, EC5, and EC6. **e** Top 20 enriched GO BP terms of specifically expressed genes in EC3, EC5, and EC6. The horizontal axis showed RichFactor (enrichment factor). Colors from blue to red represented P-value and the bubble size represented the number of genes enriched in a certain GO term. **f** A pseudotime trajectory of EC sub-clusters. The pseudotime trajectory of each EC sub-cluster was illustrated in the right. The black circles represented different cell statuses identified in the trajectory analysis. The colors indicated different samples.

To verify the sub-clustering results of ECs in normal and injured spinal cords, we further performed double immunostaining with an EC marker (CD31) and a specialized EC subtype marker, Mt2A (EC3) (Fig. 5a), Angpt2 (EC5) (Fig. 5b) and Rgs5 (EC6) (Fig. 5c), respectively. As shown in Fig. 5a, the immunostaining results were consistent with our scRNA-seq data. Also, tSNE maps of the expression distribution of these marker genes in four samples were shown in Fig. 5d.

**Fig. 5.**
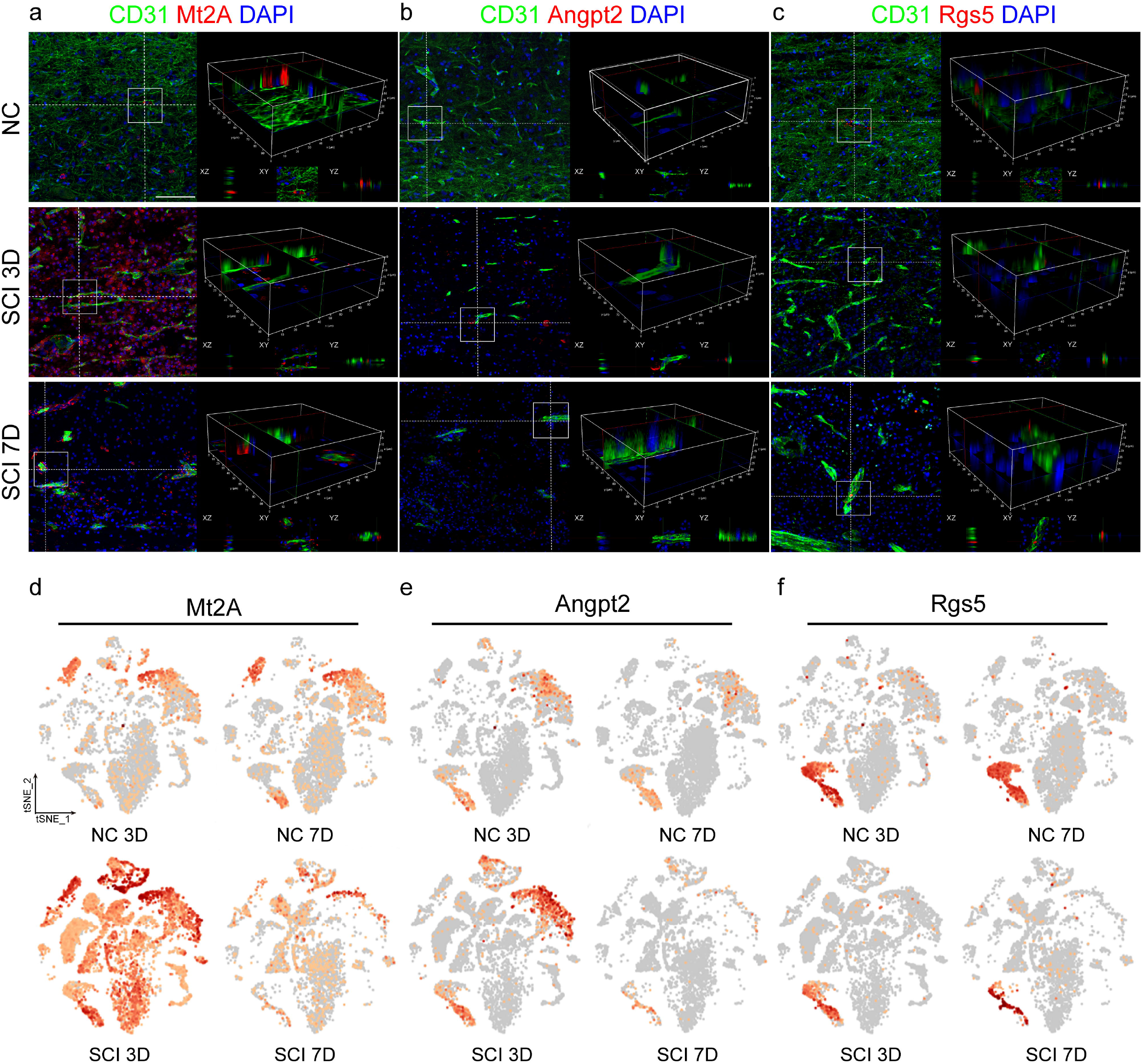
Immunohistochemistry validation of the specific ECs subtypes in the injury site. **a-c** Co-immunostaining of CD31 (green) and EC sub-cluster markers (red): Mt2A (EC3, **a**), Angpt2 (EC5, **b**), and Rgs5 (EC6, **c**), respectively, on spinal cord sections of SCI 3D, SCI 7D, and NC. Nuclei were stained with DAPI (blue). The area in the white box was selected to show in 3D view. Scale bar, 100 μm. **d-f** tSNE maps of the expression distribution of Mt2A (**d**), Angpt2 (**e**), and Rgs5 (**f**) in four samples.

### Microglia and Macrophage Regulate EC Subsets through SPP1 and IGF Signaling Pathway

We then performed Cellchat analyses to illustrate signaling pathways between EC subsets and other cell types and focused on the role of inflammation-related cells (microglia and macrophage) on ECs based on the aforementioned analysis (Fig. 3; Supplementary Table S4). The results showed that in SCI-3D and SCI-7D, microglia (C0) and macrophage (C2) regulated EC subsets through the SPP1 and IGF signaling pathways. As shown in Fig. 6a, in SCI 3D and SCI 7D, even though all kinds of cells regulate ECs through the SPP1 signaling pathway, microglia (C0) and macrophage (C2) contributed the majority interaction with ECs, especially EC4 and EC5. Also, ECs could act as ligand cells to interact with other EC sub-clusters through the SPP1 signaling pathway. Strikingly, we found that, in the IGF signaling pathway, only microglia (C0), macrophage (C2), and fibroblast (C5) participated in ECs regulation. In SCI 3D, C0, C2, C5 and EC6 regulated EC3 through the IGF signaling pathway while in SCI 7D, C0 and C2 mainly regulated EC5 (Fig. 6b). A set of ligand-receptor pairs were involved in the SPP1 and IGF signaling pathways and Fig. 6c&d demonstrated the relative contribution of each ligand-receptor pair to the overall communication network of the SPP1 and IGF signaling in SCI 3D and SCI 7D. As shown in Fig. 6c&d, Spp1-Cd44 and Igf1- (Itga6 + Itgb4) were the main L-R pairs in the SPP1 and IGF signaling, respectively. The violin plots of the main ligands and receptors involved in SPP1 and IGF signaling were shown in Fig. 6e&f. It was interesting that although Spp1-Cd44 was the main L-R pair in the SPP1 signaling, the effect on ECs was mainly through Spp1-(Itga5+Itgb1) interaction pair.

**Fig. 6.**
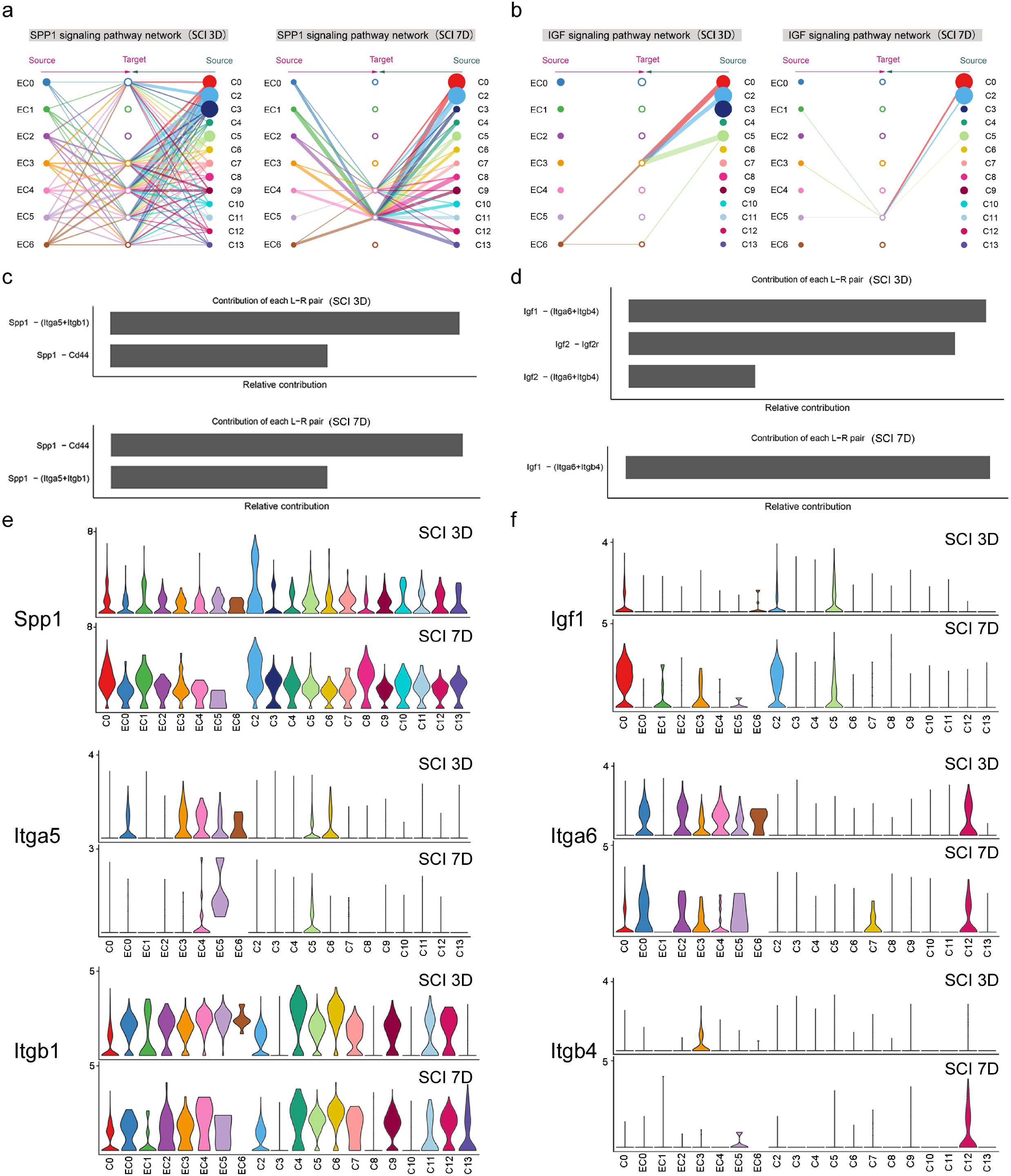
CellChat analysis of the communications between ECs subtypes and other cell types. **a**, **b** Hierarchical networks of the cell-cell communication patterns of SPP1 (**a**) and IGF signaling (**b**) when EC sub-clusters as receptors in SCI 3D and SCI 7D, respectively. Edge thickness indicated the sum of weight key signals between populations (from different cell types to EC sub-clusters). The solid circle indicated the ligand cell, and the hollow circle indicated the receptor cell type. **c**, **d** Relative contribution of each ligand-receptor pair to the overall communication network of SPP1 (**c**) and IGF signaling (**d**). **e**, **f** Violin plots of the expressions of ligands and receptors in SPP1 (**e**) and IGF signaling pathway (**f**) among each cell type in normal (NC 3D, NC 7D) and SCI (SCI 3D, SCI 7D) tissues, respectively.

## Discussion

Previously, we have performed total transcriptome sequencing on spinal cord tissues of rats at multiple time points after SCI (Yu et al., 2019). However, the whole-transcriptome sequencing has limited an in-depth analysis of cellular responses to injury due to the averaged gene expression of a group of cells. The heterogeneity of spinal cord cells make it difficult to accurately identify and reflect the heterogeneity of gene expression among different cell types. The scRNA-seq has been broadly used to address the heterogeneity, and explore the physiological and pathological functions of respective cell types in tissues. During the developmental stages of CNS and progression of several degenerative diseases, some function-related events under the regulation of the specific cell types have been deciphered by the scRNA-seq (Navin et al., 2011; Patel et al., 2014). We herein firstly identified the heterogeneity and subsets of ECs, as well as their regulatory relations with other cells in the injured rat spinal cord.

Vascular ECs compose blood vessels and play an important role in forming vascular lumen, controlling vascular permeability, and sensing cells and molecules in the circulatory system (Eilken and Adams, 2010). It was reported that the UTX gene expression increased in ECs after SCI. Tissue-specific knockdown of UTX gene in ECs promotes angiogenesis and functional recovery after SCI (Ni et al., 2019). Elevated expression of TRPV4 after SCI can damage EC structure, leading to the breakdown of the blood-spinal cord barrier (BSCB). Inhibition/knockdown of TRPV4 stabilizes vascular ECs, protects the integrity of BSCB, reduces secondary injury, and improves functional recovery after SCI (Kumar et al., 2020). These suggest that regulation of vascular ECs and vascular-related cells can promote angiogenesis and improve axonal regeneration and functional recovery after SCI.

Recently, Lindsay et al identified cellular heterogeneity and interactions with scRNA-seq after SCI in mice (Milich et al., 2021). They analyzed and divided cells at lesion sites into three groups: myeloid cells, vascular cells, and macroglial cells. ECs of the injured spinal cord were then subdivided into four subgroups in mice and the angiopoietin and VEGF signaling were identified to be important during angiogenesis (Milich et al., 2021). Another investigation by scRNA-seq after SCI pinpointed the microglia, the cells responsible for innate immunity. They analyzed FACS filtered CD45^+^ immune cells in the spinal cord from healthy and 0.5 h to 90 d post-SCI (Hakim et al., 2021), and found disease-associated microglia (DAM), which are derived from baseline microglia and are involved in functional recovery. In the present study, we clustered ECs into seven sub-clusters and mainly highlighted the interactions between inflammatory cells and ECs. Our results showed that in SCI 3D, there was a growing number of EC5, which was marked with Angpt2 and Apln. The EC5 subcluster was annotated tip cell, which was a specialized EC during sprouting angiogenesis (del Toro et al., 2010). Another interesting EC sub-cluster came from EC6, which was marked with Rgs5 and Tagln, and dramatically decreased after SCI. Rgs5 and Tagln are typical markers of mural cells (pericytes and vSMC) that are wrapped in the blood vessels (Vanlandewijck et al., 2018). We then considered that EC6 was a mural-like EC, which might mediate the interaction of EC and mural cells. The EC6 sub-cluster mainly existed in normal conditions and diminished after SCI.

ECs and immune cells are implied by previous studies that they play some roles in angiogenesis and functional recovery after SCI (Milich et al., 2021), but their interactions during angiogenesis have not be fully explored. The insights of cell communications on EC subsets indicated that microglia and macrophage interacted with ECs through SPP1 and IGF signaling pathways. SPP1, also known as osteopontin, is a phosphorylated acidic glycoprotein adhesion molecule, which has a cardiovascular function by regulating cell migration, adhesion, and spreading (Pagano and Haurani, 2006). *In vitro* evidence showed that SPP1 secreted by glioma cells regulated the proliferation, migration, and tube formation of endothelial progenitor cell (EPC), as well as promoted angiogenesis (Wang et al., 2011). Here, we demonstrated that the SPP1 was mainly secreted by macrophage and microglia to regulate ECs through Itga5 and Itgb1 receptors in the lesion after SCI. Another signal pathway identified to participate in the regulation of ECs by the inflammatory cells was IGF signaling. Our results showed that the Igf1-(Itga6+Itgb4) signaling pathway existed both in SCI 3D and SCI 7D, while Igf2-Igf2r and Igf2-(Itga6+Itgb4) signaling only appeared in SCI 3D. It is known that Igf1 is required for vessel remodeling and has a neuroprotective effect on the CNS (Lopez-Lopez et al., 2004; Ozdinler and Macklis, 2006). Our results presented that microglia, macrophage, and fibroblast were the main sources of Igf1 at the lesion site after SCI. Also, ECs (EC6 in SCI 3D, EC1, EC3, and EC5 in SCI 7D) can secret Igf1 to regulate EC3/6 at SCI 3D and EC5 at SCI 7D.

In conclusion, we first characterized the cellular heterogeneity after rat SCI and further delineated the EC subsets. We also established the interactions between ECs and other cells following SCI and showed that immune cells, like microglia and macrophage, regulated EC subsets through SPP1 and IGF signaling pathways (Fig. 7). Our results provide new insights to illustrate the complexity of EC cells and their interactions with other cells after SCI, which might facilitate the exploration of new therapeutic targets aimed at angiogenesis for SCI in the future.

**Fig. 7.**
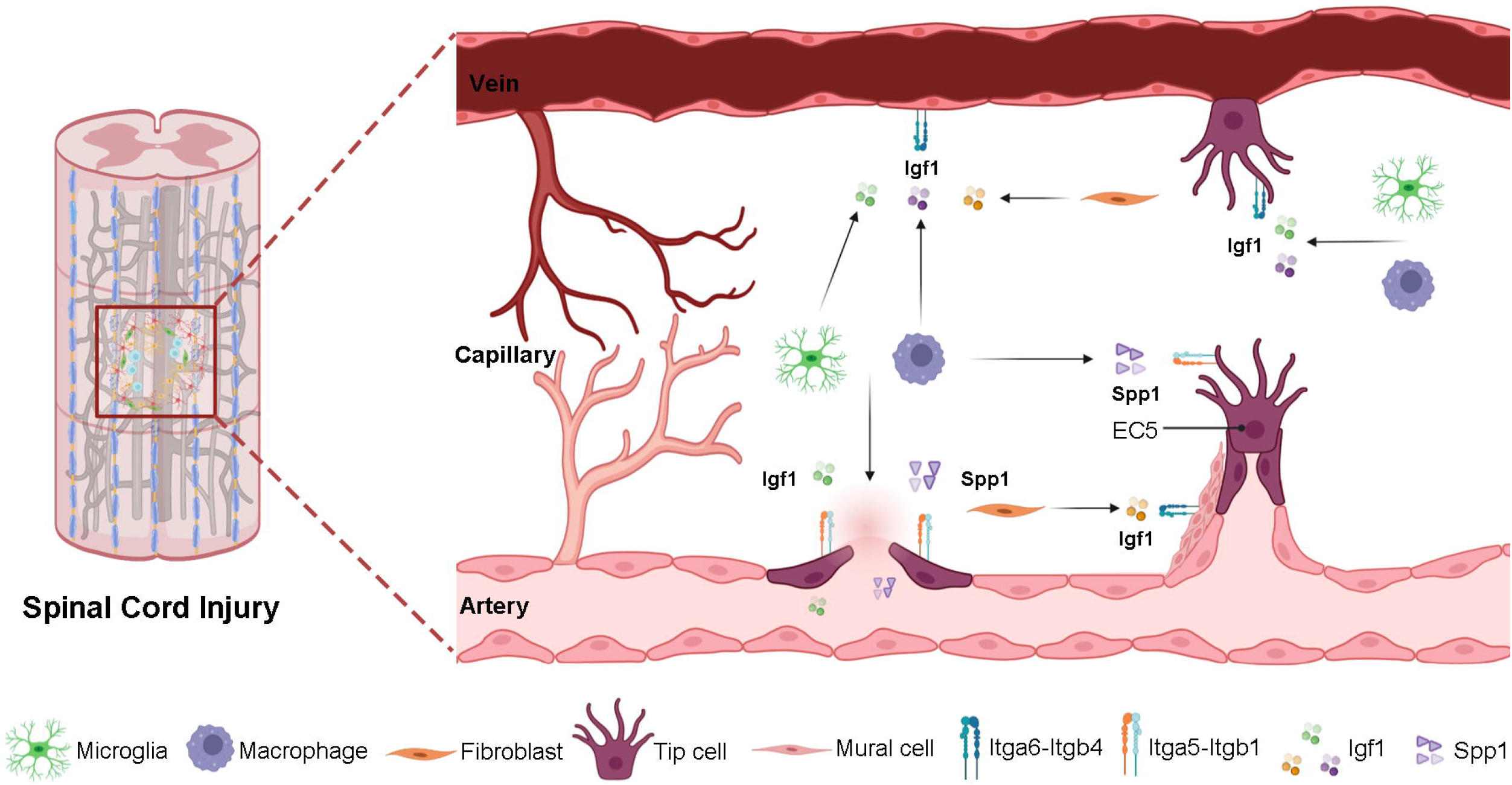
A graphic about the interactions of ECs and other cell types via SPP1 and IGF signaling after SCI. After SCI, microglia, macrophage and fibroblast secreted SPP1 and IGF1 to interact with receptors on ECs, respectively, facilitating angiogenesis.

## Materials and methods

### Experimental Design

Sprague Dawley (SD) rats underwent contusion SCI. At 3 d and 7 d post-SCI, the T9-T10 spinal cords from the injured groups and sham-operated groups were harvested. Following tissue dissociation and single cell collection, cell suspensions from six to eight rats at each group were combined into one sample for scRNA-seq. For immunostaining, spinal cord sections from injured and sham groups were stained for CD31, αSMA, GFAP, IBA1, Mt2A, ANGPT2 and RGS5 at each time point after SCI. Three animals were investigated at each time point.

### Animals and ethics statement

Female Sprague Dawley (SD) rats (210-230 g) were purchased from the Experimental Animal Center of Nantong University. All animals were maintained and used in accordance with the guidelines of Institutional Animal Care of Nantong University. All animal experiments were approved by the Institutional Animal Ethics Committee of Nantong University, China. Animals were housed in temperature-controlled (24°C) and humidity-controlled (50%) animal quarters with a 12 h light/dark cycle and had free access to food and water.

### Animal surgery models

All rats underwent a surgical procedure as previously described (Zhu et al., 2015). Rats were randomly divided into four groups: SCI-3D, SCI-7D, NC-3D and NC-7D. Before surgery, the rats were anesthetized with an injection of pentobarbital sodium (40 mg/kg, intraperitoneal [i.p.]) and received a T10 contusion injury. Laminectomy was then performed aseptically at T9-T10 to expose the spinal cord followed by stabilization via a spinal frame clamping the T9-T11 spinous processes. The contusion SCI was made by Infinite Horizon Impactor device (Precision Systems and In-strumentation, LLC) at 160 kdyn for 30 s. Animals in the sham group only received a laminectomy. Post-surgery, bladder dysfunction was present in all SCI animals and bladders were manually emptied twice a day for the duration of the experiments.

### Immunohistochemistry

The collected injury epicenter spinal cords were fixed in 4% paraformaldehyde (PFA) overnight, and dehydrated in 30% sucrose solution for 3 d at 4°C. Then, the tissues were embedded in Optimal Cutting Temperature compound (OCT, Sakura) and sectioned into 30-μm slices using a cryostat (Leica CM3050 S). Frozen tissues stored at −80°C. For immunohistochemistry, tissue sections were incubated with indicated primary antibodies overnight at 4°C: CD31 (R&D, AF3628, 1:200), GFAP (Abcam, ab4674, 1:500), IBA1 (wako, 019-19741, 1:500), αSMA (Cell Signaling Technologies, 19245S, 1:500), Mt2A (Thermo Fisher Scientific, MA1-25479, 1:500), ANGPT2 (Proteintech, 24613-1-AP, 1:100), RGS5 (Novus, NBP2-00880, 1:150). The next day, the sections were incubated in blocking solution containing secondary antibody at 1:500 dilution (donkey-anti-goat Alexafluor plus-488, donkey-anti-rabbit Alexafluor-568, donkey-anti-mouse Alexafluor-594, donkey-anti-chicken Alexafluor-647) for 2 h at room temperature. Images were acquired with a Leica DMi8 confocal microscope and processed with Leica Microsystems imaging software (LAS X, version 4.1).

### Single cell collection for spinal cord tissues

Spinal cord tissues were minced and enzymatically digested with collagenase I (Sigma) and 0.5 mg/ml DNase I (Sigma) for 10 min at 37°C with shaking. Then, DMEM containing 10% FBS was added to the samples to stop the digestion. After filtering through a 70-μm cell strainer, the cell suspensions were pelleted at 300 ×g for 10 min at 4°C, and re-suspended in PBS with 1% BSA. Red blood cell lysis buffer was added and incubated for 2 min to lyse red blood cells. The single cell suspensions were re-filtered through a 30-μm strainer (Corning) and further enriched with a dead cell removal kit (Miltenyi Biotec) according to the manufacturer’s protocol.

### Single-cell RNA-seq library preparation and sequencing

Cellular suspensions were loaded on a 10X Genomics GemCode Single-cell instrument that generates single-cell Gel Bead-In-EMlusion (GEMs). Libraries were generated and sequenced from the cDNAs with Chromium Next GEM Single Cell 3’ Reagent Kits v3.1. Libraries were then pooled and sequenced on an Illumina NovaSeq 6000. The raw data were converted to FASTQ files, underwent alignment and counts quantification with 10X Genomice Cell Ranger software (10x Genomics, version 3.1.0) (Gene Denovo, Guangzhou, China)).

### Single-cell RNA-Seq data processing

The cells by gene matrices for each sample were individually imported to Seurat (version 3.1.1) for downstream analysis. Qualified cells with mitochondrial gene percentage less than 20% and detected gene number > 200 were kept. Data were then log-normalized for the subsequent analyses.

### Dimension reduction and cell clustering

We performed principal component analysis (PCA) dimensionality reduction with the highly variable genes as input in the Seurat package. The first 20 principal components of PCA were used to cluster the cells. The total clustering analysis was performed with a graph-based clustering approach implanted in Seurat, based on the shared nearest neighbor (SNN) clustering. For sub-clustering analysis, the ECs were extracted and clustered in a similar procedure. Following clustering, t-distributed stochastic neighbor embedding (t-SNE) was generated for visualization of clusters.

### Cluster Annotation

To annotate cell clusters, we applied the FindAllMarkers () function in Seurat using the default non-parametric Wilcoxon rank sum test with Bonferroni correction. The cell groups were annotated based on the DEGs and the well-known cellular markers from the literature.

### Differentially expressed genes analysis and Gene Ontology (GO) enrichment analysis

The specific genes of each cluster were analyzed by wilcoxon algorithm (Ntranos et al., 2019) and scored by group one vs rest. Genes with high cluster-specific expression, logFC > 0.25 and expressed in at least 20 % of cells were selected as differentially expressed genes. Differentially expressed genes in different cell cluster or different samples were subjected to Gene ontology (GO) analysis (Ashburner et al., 2000). Significantly enriched GO terms were identified using Fisher’s exact test.

### Pseudotime Analysis for ECs

Pseudotime analysis was conducted for endothelial cells. We employed Monocle 2 to illustrate the pseudotime trajectories that projected the high-dimensional transcriptomic data to one dimension.

### Cell-cell communication analysis

We applied CellPhoneDB method to analyze potential cell-cell interactions between different cell types (Vento-Tormo et al., 2018). Only receptors and ligands expressed more than 10% of the cells were considered for analysis in specific clusters. Interaction was constructed as a receptor-ligand pairing matrix. Significant interactions can then be prioritized on the basis of their P value. To further identify signaling pathways involved in the interactions between EC and other cell types, we used CellChat (v1.1.0; https://github.com/sqjin/CellChat) R package to predict major signaling inputs and outputs of cells and how these cells and signals coordinate for functions.

## Acknowledgements

This work was supported by the National Key R&D Program of China (No. 2020YFA0113600), the National Natural Science Foundation of China (No. 32130060) and the Natural Science Foundation of Jiangsu Province (No. BK20202013). We thank Shanghai Biotechnology Corporation and Genedenovo Biotechnology Co., Ltd for assisting in sequencing and analyzing.

## Conflict of interests

The authors declare no competing financial interests.

## Author contributions

B.Y., Y.J.W. and C.Y. conceived and designed the experiments. Y.Q.C., Y.H.L. and D.W. performed the experiments. C.Y. and Y.Q.C. analyzed the data. C.Y., Y.Q.C and Y.J.W. wrote the manuscript. B.Y., X.H.W., Y.J.W., Y.L. and X.S.G revised the manuscript. All authors have read and approved the final manuscript.

---

**Supplementary Fig. S1 Immunostaining of vSMCs and astrocyte in the injured spinal cord. a** Co-immunostaining of CD31^+^ ECs (green) and αSMA^+^ vSMCs (red). Arrows indicate the interacted ECs and vSMCs. **b** Co-immunostaining of CD31^+^ECs (green) and GFAP^+^ astrocyte (red). Arrows indicate the interacted ECs and astrocytes. The three small squares below were the enlarged images of the white square in each rectangle image. The border between uninjured and injured was drawn with the dashed line. Nuclei were stained with DAPI (blue). Scale bar: 500 μm and 100 μm, respectively.

**Supplementary Fig. S2 Characteristics and sampling metrics for single-cell sequencing.** Detected gene numbers and total UMIs of each cluster. **a-b**, The box plot showed the distribution of detected gene numbers (A) and total UMIs per cell (B) of single cells in each of the 26 clusters. **c**, tSNE cluster map showed the distribution of all cells in 26 clusters.

**Supplementary Fig. S3 Cell-type-specific interactions between ECs and the rest of the cell types when EC was the ligand cell.** Circle sizes represented significance, which were defined as −log10 (p value+1e-04). Colors from green to red represented the interaction score between ligands and receptors.

**Supplementary Fig. S4 Gene expression and GO analysis heterogeneity of ECs sub-clusters. a** The expression heatmap of DEGs among ECs sub-clusters. **b**. Violin plots of the expression levels of marker genes in EC0 (Car4), EC1 (Hba-a2.1), EC2 (Adtrp), and EC4 (Serpina1). **c**. Enriched top 20 enriched GO BP of DEGs in EC0, EC1, EC2, and EC4.

## Notes

### Competing Interest Statement

The authors have declared no competing interest.

