## Supplementary figures and images for "Single cell sequencing reveals microglia induced angiogenesis by specific subsets of endothelial cells following spinal cord injury"

### Supplementary FigS1

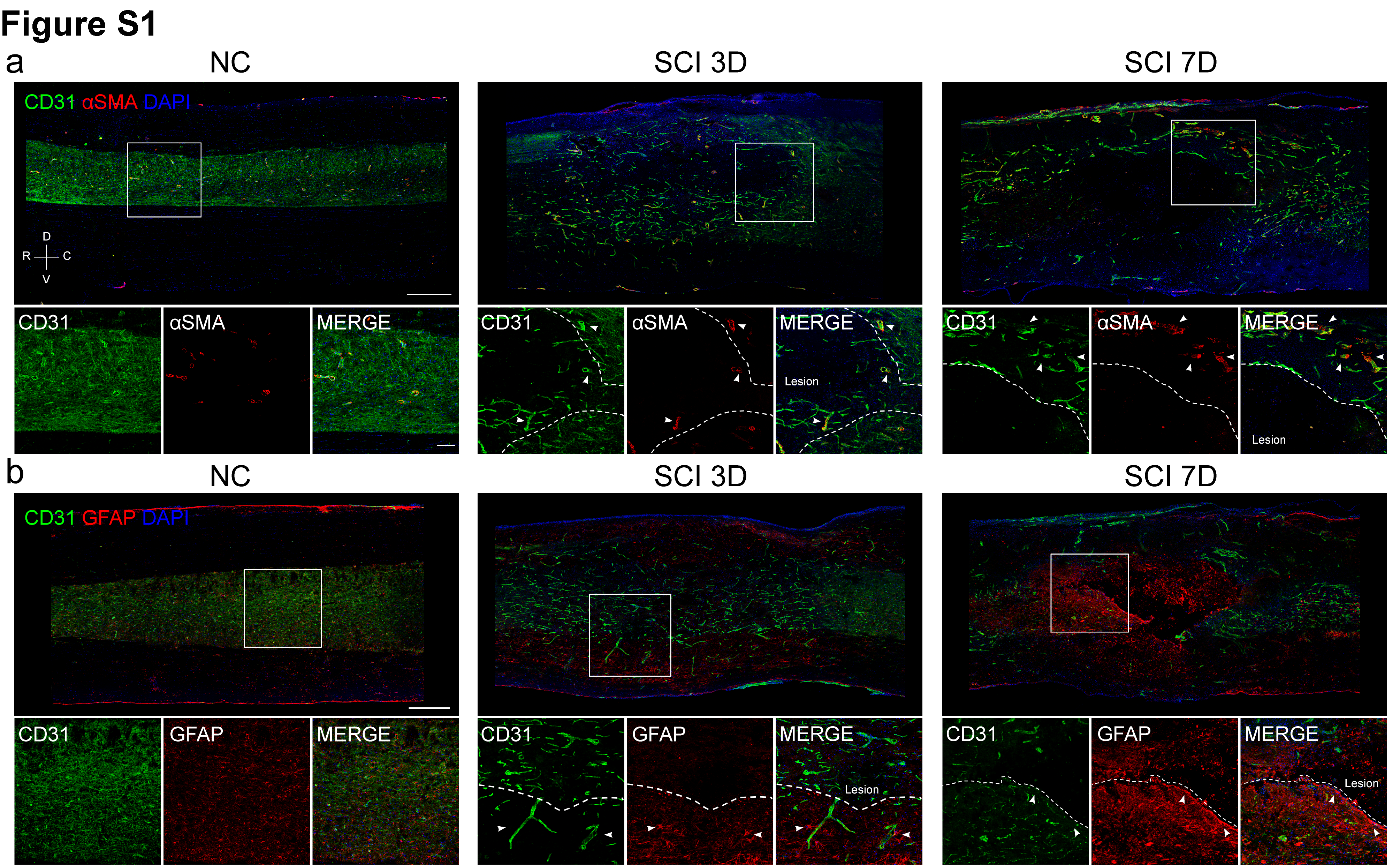

### Supplementary FigS2

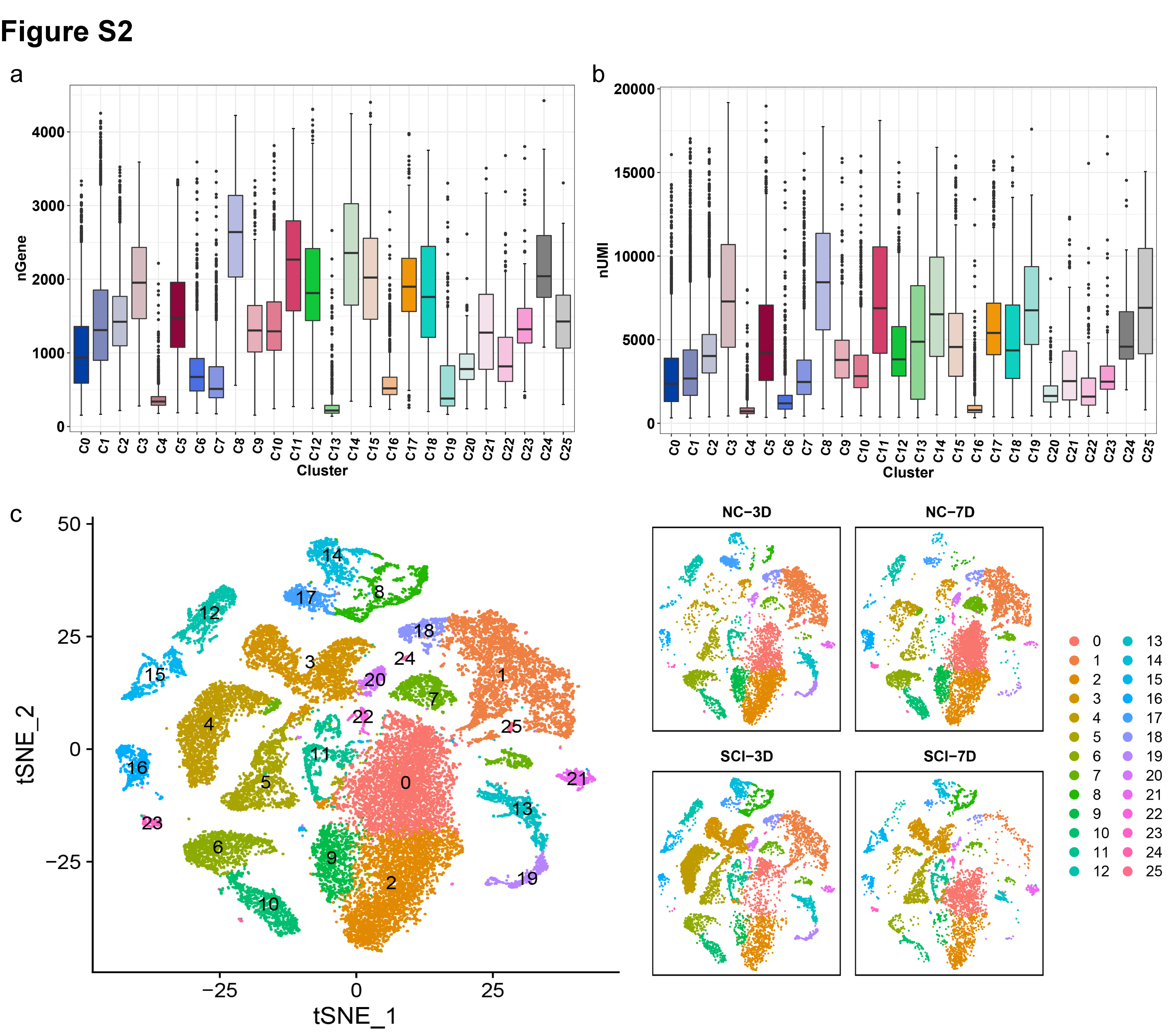

### Supplementary FigS3

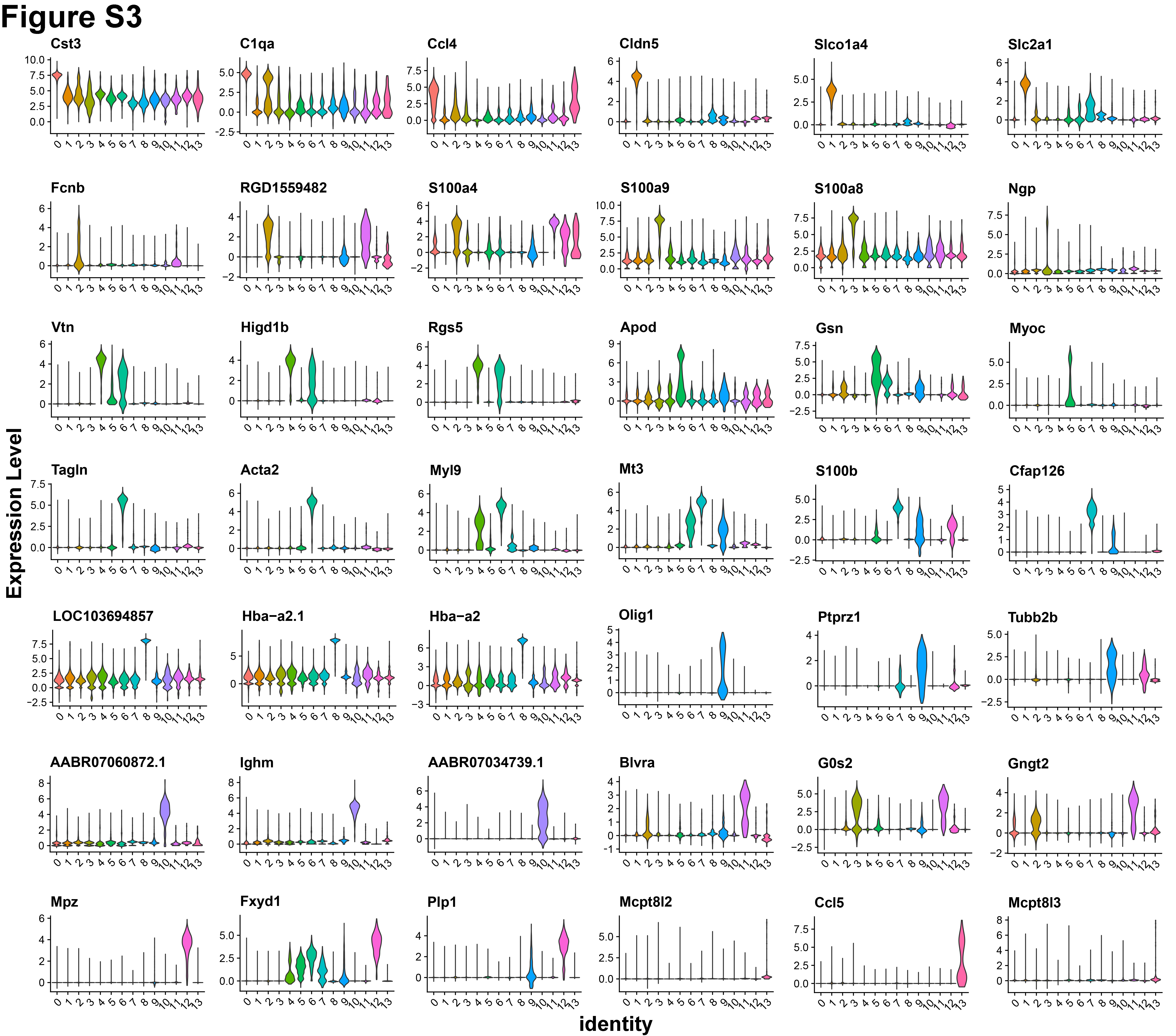

### Supplementary FigS4

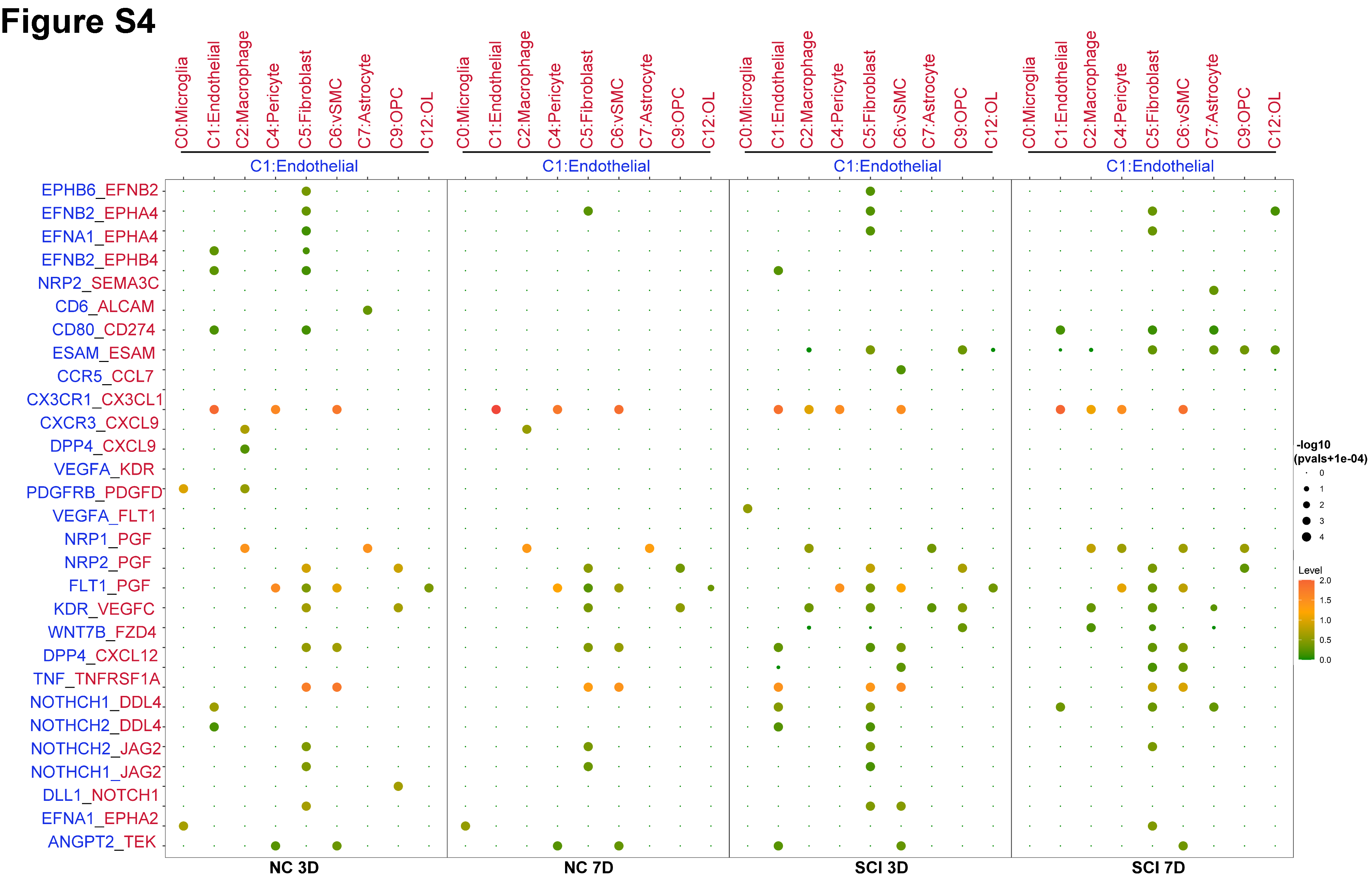

### Supplementary FigS5

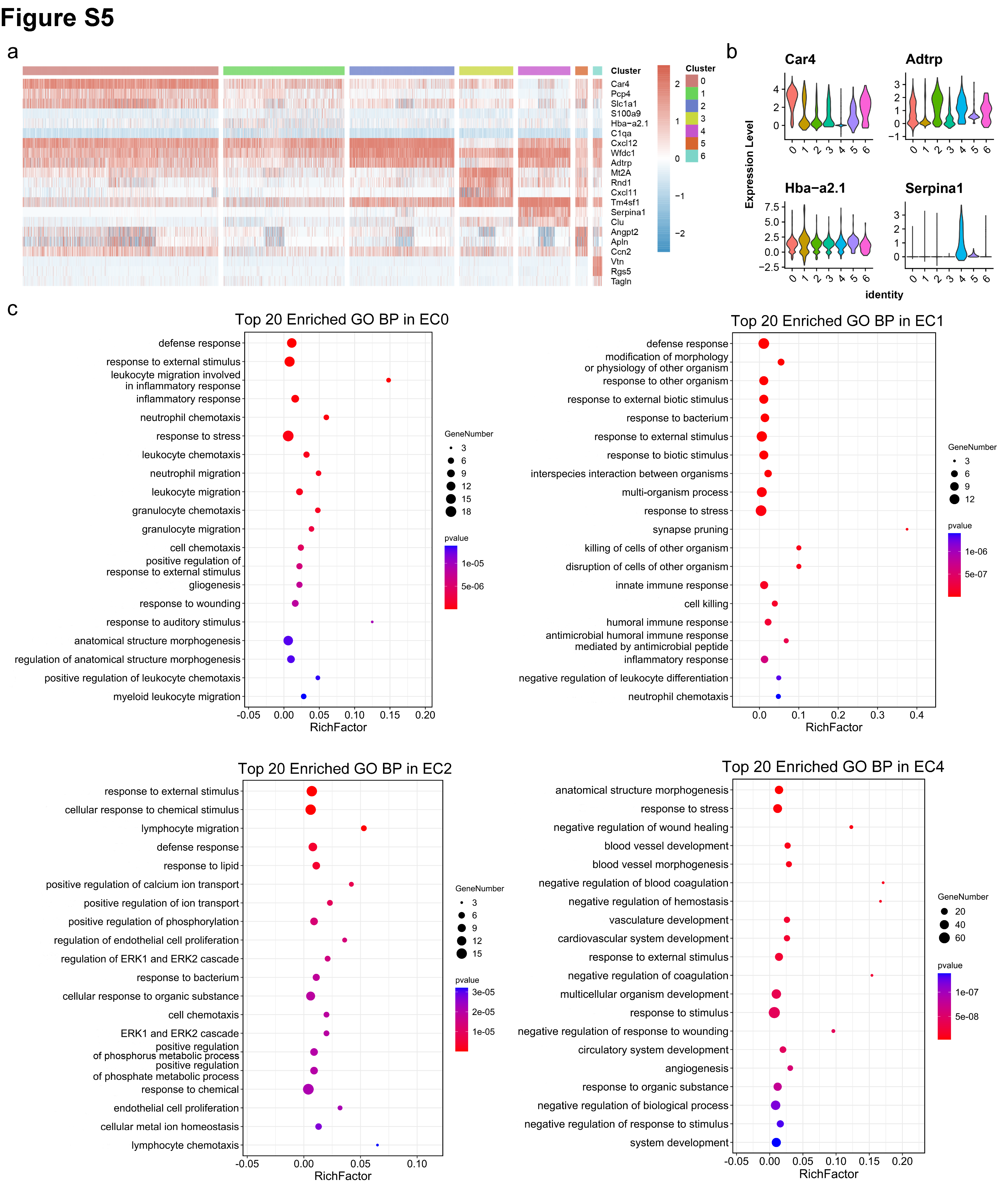
